# Efficient Heuristic for Decomposing a Flow with Minimum Number of Paths

**DOI:** 10.1101/087759

**Authors:** Mingfu Shao, Carl Kingsford

## Abstract

Motivated by transcript assembly and multiple genome assembly problems, in this paper, we study the following *minimum path flow decomposition* problem: given a directed acyclic graph *G* = (*V,E*) with source *s* and sink *t* and a flow *f*, compute a set of *s-t* paths *P* and assign weight *w*(*p*) for *p* ∈ *P* such that 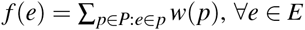, and |*P*| is minimized. We propose an efficient pseudo-polynomial-time heuristic for this problem based on novel insights. Our heuristic gives a framework that consists of several components, providing a roadmap for continuing development of better heuristics. Through experimental studies on both simulated and transcript assembly instances, we show that our algorithm significantly improves the previous state-of-the-art algorithm. Implementation of our algorithm is available at https://github.com/Kingsford-Group/catfish.

## 1 Introduction

RNA sequencing (RNA-seq) is an established technology that enables identification of novel genes and transcripts as well as accurate measurement of expression abundances [1]. The RNA-seq protocol produces short sequencing reads sampled from the expressed transcripts. Thus, a fundamental computational problem is to recover the set of full-length expressed transcripts and their expression abundances in the sample from these reads. This is referred to the *transcript assembly* problem. There have been much research interests for this problem and many methods and software packages have been proposed. Depending on whether a reference genome is assumed available, these methods are divided into two categories, *reference-based* methods (e.g., Cufflinks [2], Scripture [3], IsoLasso [4], SLIDE [5], CLIIQ [6], CEM [7], MITIE [8], iReckon [9], Traph [10], and StringTie [11]), and *de novo* methods (e.g., TransABySS [12], Rnnotator [13], Trinity [14], SOAPdenovo-Trans [15], Velvet [16], and Oases [17]). However, according to the bench-marking studies [18, 19], the performance of these methods are far from satisfactory, especially when the expressed genes contain multiple splice isoforms. Hence, transcript assembly still remains an open and challenging problem, and thus requires further algorithm development.

Both reference-based and *de novo* methods share the similar pipeline. The first step of the pipeline is to build the so-called *splice graph* (or transcript graph, overlap graph, etc) from given reads. For reference-based methods, reads are first aligned to the reference genome using some spliced aligner (e.g., TopHat2 [20], STAR [21], and HISAT [22]). Then the boundaries of exons and introns are inferred from the spliced aligned reads, and the splice graph is consequently constructed: each vertex represents an exon (or partial exon), each edge indicates there exist reads spanning the two corresponding exons, and the weight of this edge is usually computed as the number of such spanning reads. For *de novo* methods, reads are usually first organized by the *de bruijn* graph. Then the splice graph is constructed by collapsing simple paths of the *de bruijn* graph, and the weight of each edge is computed as the number of reads associated with the corresponding path. For both categories of methods, the constructed splice graph is the superposition of the expressed transcripts, each of which corresponds to an (unknown) path in the splice graph. If we assume the ideal case that the read coverage for each expressed transcript is a constant across the whole transcript, then the weights of edges shall form a flow of the spliced graph. In the realistic scenario where the read coverage varies across each transcript, there exist methods that can smooth the edge weights to make it a flow [10].

The second step of the pipeline is usually to decompose the splice graph into a set of paths and associate a weight with each path. These predict the expressed transcripts and their expression abundances. This step is the most algorithmically challenging part in the whole transcript assembly pipeline. Under the principle of parsimony, it is usually formulated as an optimization problem of decomposing a given flow into a minimum number of paths. Formally, let *G* = (*V*, *E*) be a directed acyclic graph (DAG) with a source vertex *s* and a sink vertex *t*. We denote the set of all *s*-*t* paths of *G* by 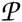. Notice that 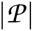 might be exponential compared with |*E*|. Let *f*: *E* → *ℤ* be an *s*-*t* integral flow of *G*. We say (*P*, *w*) is a *decomposition* of (*G*, *f*) if we have 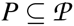, *w*: *P* → *ℤ* assigns an integral weight *w*(*p*) for each path *p* ∈ *P*, and *f* (*e*) = ∑ _*p*∈*P*:*e*∈*p*_ *w*(*p*) for every *e* ∈ *E*. Our paper focuses on the following problem.

### Problem 1 (Minimum Path Flow Decomposition)

*Given a DAG G* = (*V, E*) *with source vertex s and sink vertex t and an integral s-t flow f, compute a decomposition* (*P*, *w*) *of* (*G*, *f*) *such that* |*P*| *is minimized.*

While we formulate Problem 1 in the context of transcript assembly, we emphasize that Problem 1 can be naturally used to model a broader class of so-called *multiple genome* assembly problems [23]. These are problems in genomics where several unknown strings *S* = {*s*_1_, *s*_2_, …, *s*_*k*_} and their (unknown) counts *C* = {*c*_1_, *c*_2_, …, *c*_*k*_} must be reconstructed from many substrings (reads) that are sampled from these strings. This is a generalization of the standard genome assembly problem that seeks to reconstruct a single string. The above-mentioned transcript assembly problem is one such instance, where *S* is the set of alternatively spliced isoforms of a gene, and *C* is the expression levels of these isoforms. Other instances include metagenomic assembly (where *S* represents the genomes of microbes in a community and *C* represents their abundances), cancer genome assembly (where *S* is the set of genomes of subclones in a tumor sample and *C* is the frequency of each subclone), and virus quasi-species assembly (where *S* is the set of genomes in a population of viral genomes infecting a given individual and *C* is their frequency).

Problem 1 has been proven to be strongly NP-hard [24], meaning that even if all the flow values are in polynomial-size, there still does not exist a polynomial-time algorithm unless P = NP. It has been further proved that even if all flow values are chosen from {1, 2, 4}, this problem is still NP-hard [25].

Several algorithms and heuristics have been proposed for Problem 1. Vatinlen et al. [24] proposed two practical greedy algorithms, namely *greedy-length* and *greedy-width* algorithms, which are to iteratively choose the shortest path and the path with the largest flow, respectively, in the remaining flow, until the entire flow is decomposed. These two algorithms have been shown that they can guarantee obtaining a decomposition with at most (|*E*| − |*V*| + 2) paths, a known upper bound for the number of paths in any optimal solution. Hartman et al. [25] designed two similar algorithms, called *bicriteria-width* and *bicriteria-length*, which can guarantee decomposing (1 − *ε*) fraction of the entire flow into *O*(|*P*^∗^|/*ε*^2^) paths, for any ε < 1. Mumey et al. [26] proposed another two algorithms using parity balancing path flows, which have been proved to be (*L*^log(*F*)^ · log(*F*))-approximation algorithms, where *L* is the maximum length of all *s*-*t* paths in *G*, and *F* is the maximum flow values over all edges. We emphasize that over all existing algorithms, the greedy-width algorithm has been shown as the best heuristic through experimental comparisons [24, 25, 26], and because of that, this algorithm has been implemented in several widely used transcript assembly methods, for example, Traph [10] and StringTie [11].

Several problems related to Problem 1 have been also addressed. Hendel and Kubiak [27] studied the problem of decomposing a given flow into a set of paths such that the length of the longest path is minimized. They proposed a polynomial-time approximation scheme (PTAS) for this problem. Baier et al. [28, 29] studied the *k-splittable flow* problem, which is to compute a maximum flow such that it can be decomposed into at most *k* paths. For this problem they devised a (2/3)-approximation algorithm for *k* = 2 and a (2/k)-approximation algorithm for *k* ≥ 3.

Our contribution to Problem 1 is a pseudo-polynomial-time heuristic based on the novel properties we prove, and we show that it considerably outperforms the best previous heuristic (greedy-width) through extensive experimental comparisons. The central idea of our heuristic is that equations involving flow values on edges are intuitively suggestive of the structure of paths. For example, if *f* (*e*_1_) = *f* (*e*_2_) + *f* (*e*_3_), then one might suspect that a path goes through *e*_1_ and *e*_2_ and another path goes through *e*_1_ and *e*_3_ (see examples in Figure 1). The first challenge of our heuristic is to identify these equations. Once identified, each equation can then be used to simplify the graph by merging edges that are involved in the same path. We define a operation, called reverse, on the graph that modifies the graph in such a way that the optimal solution does not change, but that attempts to make edges that are to be merged be incident on the same vertex. The entire approach iterates between these two steps: identifying a non-trivial relationship between the flow values on the edges and modifying the graph to facilitate edge merging (and merging the edges). We show that at each stage, either the algorithm fails, or the solution space gets reduced and the gap between the optimal and an upper bound on the optimal is reduced.

**Figure 1:**
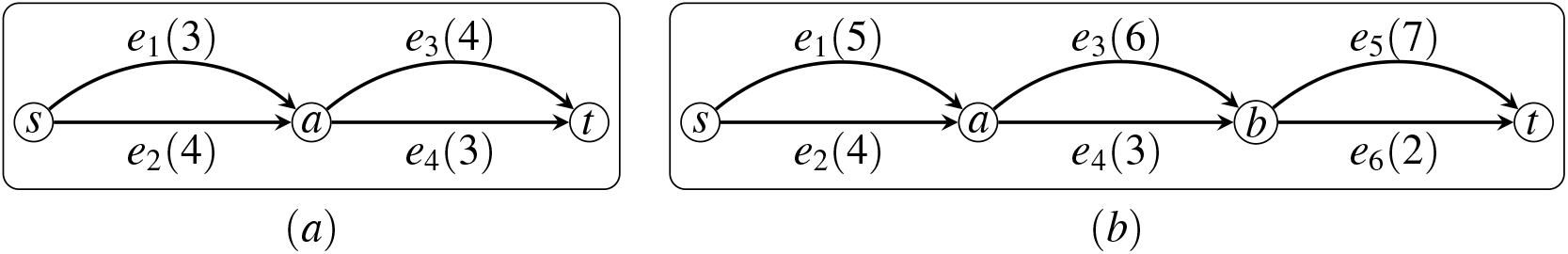
The flow value for each edge is put in parenthesis. **(a)** Observe that *f* (*e*_1_) = *f* (*e*_4_), i.e, nontrivial null vector of (1, 0, 0, −1)^*T*^. We have Δ = 3, 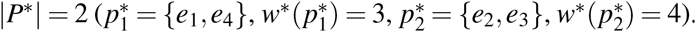. **(b)** Observe that *f* (*e*_1_) = *f* (*e*_4_) + *f* (*e*_6_), i.e., nontrivial null vector of (1, 0, 0, −1, 0, −1)^*T*^. We have **Δ** = 4, 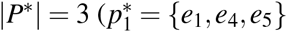, 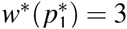, 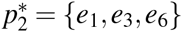, 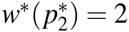, 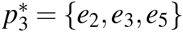, 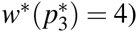

## 2 Motivation

In this Section, we prove some properties about Problem 1 upon which we design our algorithm. We first reformulate some definitions to facilitate using algebra tools. We arbitrarily assign indices to edges in *E* and paths in 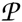, i.e., *E* = {*e*_1_, *e*_2_, …, *e*_|*E* |_} and 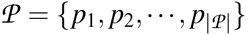. We can then write *f* as a row vector in *ℤ*^1×|*E*^ ^|^, with *f* [*k*] = *f* (*e*_*k*_). We can also write 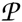 as a matrix in {0, 1}^|*P*^ ^|×|*E*^ ^|^, with 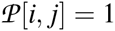 if and only if *p*_*i*_ contains edge *e*_*j*_. Consequently, each subset 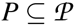 can also be represented as a binary matrix, which can be modified from matrix 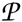 by removing these rows whose corresponding paths are not in *P*. With these formulations, the definition that (*P*, *w*) is a decomposition of (*G*, *f*) can be equivalently written as *f* = *w* · *P*, where *w* ∈ *ℤ*^1×|*P*|^. In the following, we use 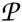 (and *P*) to represent both the set of paths as well as the corresponding matrix, but this will be obvious to distinguish. Let (*P*^∗^, *w*^∗^) be an optimal decomposition of (*G*, *f*). Let Δ= |*E*| − |*V*| + 2. From [24], we know that Δ gives an upper bound for |*P*^∗^|, i.e., |*P*^∗^| ≤ Δ. We call (Δ − |*P*^∗^|) as the *optimality gap*. Let 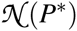 be the null space of *P*^∗^, i.e., 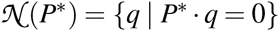. We call vectors in 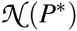 as *null vectors* (*w.r.t*. *P*^∗^). The proofs of all Propositions are in the Appendix C.

### Proposition 1

*We have f* · *q* = *0 for any* 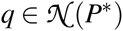.

Proposition 1 is a necessary condition rather than a sufficient condition to characterize 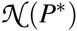. On the other side, if we assume that all weights in *w*^∗^ are randomly assigned, then it is with small probability to observe that *f* · *q* = 0 if 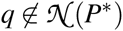.

### Proposition 2

*Suppose* 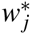 *is sampled independently and uniformly at random from* ℤ_*k*_ = {1, 2, …, *k*} *for all j* = 1, 2, …, |*P*^∗^|. *Then we have* 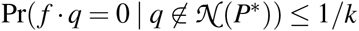.

We further study the structure of 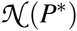. Let *V*_0_ = *V* \ {*s*,*t*}. For each *v* ∈ *V*_0_, we define a column vector *q*_*v*_ ∈ {−1, 0, +1}^|*E*^ ^|×1^ where *q*_*v*_[*k*] = 0 if *e*_*k*_ is not adjacent to *v*, *q*_*v*_[*k*] = +1 if *e*_*k*_ points to *v*, and *q*_*v*_[*k*] = −1 if *e*_*k*_ leaves *v*. Let 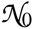 be the linear space spanned by {*q*_*v*_ | *v* ∈ *V*_0_}. We call vectors in 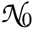 as *trivial null vectors*, and call vectors in 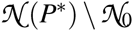 as *nontrivial null vectors*. We have the following results.

### Proposition 3

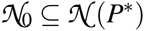.

### Proposition 4

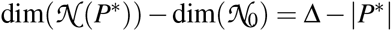.

Proposition 4 says that the optimality gap is larger than 0 if and only if there exists nontrivial null vectors (see Figure 1 for examples). This inspires our algorithm: suppose that we have two oracles, the first one can return nontrivial null vectors, and the second one can use these vectors to reduce the optimality gap; then we can iteratively apply these two oracles, and when the first oracle returns the empty set, the optimality gap has been closed. Our algorithm then tries to approximate these two oracles.

## 3 Algorithm

Our heuristic for Problem 1 consists of two phases. In its first phase, in each iteration, it tries to identify nontrivial null vectors (see Section 3.1), and then use them to partially decompose and update the graph and flow (see Section 3.2) in order to obtain a new graph with a smaller optimality gap. We expect that, the optimality gap can be reduced by at least 1 after each iteration, and the entire optimality gap can be closed after the first phase. Its second phase uses the greedy-width algorithm to decompose the remaining graph, which guarantees optimality if the optimality gap can be correctly closed in the first phase. The entire algorithm is formally described and analyzed in Section 3.4.

### 3.1 Identifying Nontrivial Null Vectors

The idea to identify nontrivial null vectors is based on Proposition 1 and Proposition 2, i.e., to compute vector 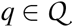 satisfying that *f* · *q* = 0 and that 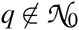, where 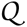 is a collection of candidate vectors that we guess in advance. We say a column vector *q* ∈ ℝ^|*E*^ ^|×1^ is *simple* if all elements of *q* are in {−1, 0, +1}. We choose 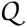 as the set of all simple vectors. Notice that it could be the case that nontrivial null vectors exist, but none of them is simple (Figure 2). Let 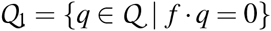. For any subset *E*_1_ ⊆ *E*, we define *f* (*E*1) = 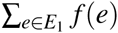. We say two subsets *E*1, *E*2 ⊆ *E* are balanced if *f* (*E*1) = *f* (*E*2). For a vector 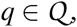, we define *E*_*s*_(*q*) = {*e*_*j*_ ∈ *E* | *q*[ *j*] = +1}, and define *E*_*t*_ (*q*) = {*e*_*j*_ ∈ *E* | *q*[ *j*] = −1}. It is clear that for any 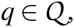, we have 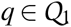 if and only if *f* (*E*_*s*_(*q*)) = *f* (*E*_*t*_ (*q*)). We say a pair of balanced subsets *E*_1_ and *E*_2_ are *indivisible*, if there does not exist two strict subsets 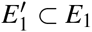 and 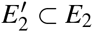 such that 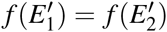. Notice that two balanced subsets being indivisible implies that they are disjoint. We also say 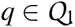 is *indivisible* if *E*_*s*_(*q*) and *E*_*t*_ (*q*) are indivisible. Let 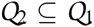 be the subset that includes all indivisible vectors. We focus on computing indivisible and nontrivial null vectors, formally illustrated as Problem 2. We emphasize that 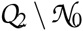 is an approximation of 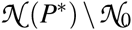, i.e., vectors in 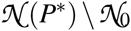 are not necessarily in 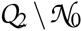 (because we restrict to simple vectors, see Figure 2), and vectors in 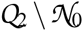 are not necessarily in 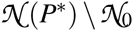 (because *f* · *q* = 0 is not a sufficient condition).

**Figure 2:**
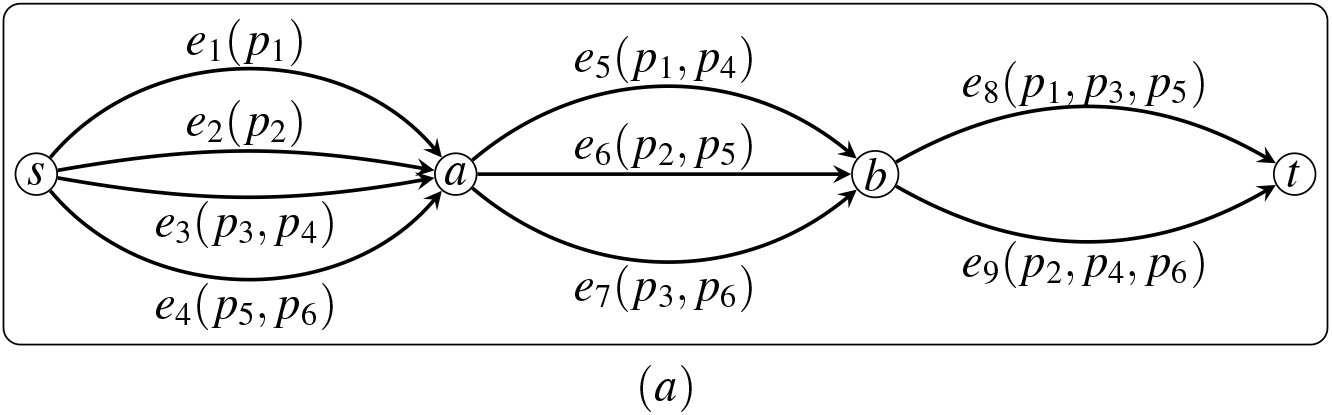
In this example, *P*^∗^ contains six paths, {*p*_1_, *p*_2_, …, *p*_6_}. The set of paths that go through each edge are labeled in the parenthesis. We also have Δ= 7, thus there exists nontrivial null vectors, for example, (2, −1, 1, 0, −1, 1, 0, −1, 0)^*T*^. However, we can verify there does not exist simple ones for this example.

#### Problem 2

*Given G and f*, *compute* 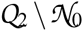.

We give a heuristic for Problem 2. We first devise a dynamic programming heuristic to compute 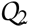 (Algorithm 1), which extends the pseudo-polynomial-time algorithm for the subset-sum problem, then we devise an algorithm (Algorithm 2) based on the properties of 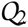 and 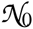 to filter out vectors in 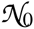.

#### Algorithm 1 Heuristic to compute 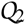

**Input:** *G* and *f* (we define | *f* | = ∑_*e*∈*E*:*e*=(*s*,*v*)_ *f* (*e*) = ∑_*e*∈*E*:*e*=(*v*,*t*)_ *f* (*e*), i.e., the value of *f*)

**Output:** 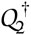

1. Let *A* ∈ {0, 1}^|*E*^ ^|×|^ ^*f*|^: *A*[*k*, *x*] = 1 if there exists *E*_1_ ⊆ {*e*_1_, …, *e*_*k*_} with *f* (*E*_1_) = *x*.

2. Let *B* ∈ {0, 1}^|*E*^ ^|×|^ ^*f*^ ^|^: *B*[*k*, *x*] = 1 if there exists distinct *E*_1_, *E*_2_ ⊆ {*e*_1_, …, *e*_*k*_} with *f* (*E*_1_) = *f* (*E*_2_) = *x*.

3. Compute *A* with the recursion: *A*[*k*, *x*] = 1 if and only if *A*[*k* − 1, *x*] = 1, or *A*[*k* − 1, *x* − *f* (*e*_*k*_)] = 1.

4. Compute *B* with the recursion: *B*[*k*, *x*] = 1 if and only if *B*[*k* − 1, *x*] = 1, or *B*[*k* − 1, *x* − *f* (*e*_*k*_)] = 1, or *A*[*k* − 1, *x*] = 1 and *A*[*k* − 1, *x* − *f* (*e*_*k*_)] = 1.

5. Compute *C* = {1 ≤ *x* ≤ | *f* | | *B*[|*E* |, *x*] = 1}.

6. Compute *D* = {*x* ∈ *C* | *B*[|*E* |, *x* − *f* (*e*)] = 0, ∀*e* ∈ *E*}.

7. For every *x* ∈ *D*, retrieve the two balanced subsets through tracing back matrices *A* and *B*, and then add the corresponding vector to 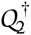.

By the definition of matrix *B*, for each *x* ∈ *C*, there exists two subsets *E*_1_ and *E*_2_ such that *f* (*E*_1_) = *f* (*E*_2_) = *x*, but these two subsets might be not indivisible (or even not disjoint). We can prove that the two balanced subsets through tracing back from *x* ∈ *D* are guaranteed indivisible (see proof of Proposition 5).

#### Proposition 5

*We have that* 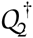, *i.e., the set returned by Algorithm 1, is a subset of* 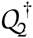. *Algorithm 1 runs in O*(|*E*| · |*f*|) *time*.

The remaining task is to compute 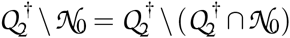. The following proposition gives an equivalent condition to decide whether a vector 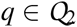 is in 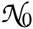 or not. We say a subset *V*_1_ ⊆ *V* is *connected* if the subgraph of *G* induced by *V*_1_ is weakly connected.

#### Proposition 6

*Let* 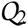. *We have that* 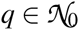 *if and only if there exists a connected subset V*_1_ ⊆ *V*_0_ *such that* 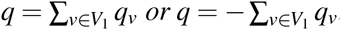.

For a subset *V*_1_ ⊆ *V*_0_, we define *E*_*s*_(*V*_1_) and *E*_*t*_ (*V*_1_) the sets of edges corresponding to cut (*V*_1_,*V* \ *V*_1_), i.e., *E*_*s*_(*V*_1_) = {(*u*, *v*) ∈ *E* | *u* ∈ *V* \ *V*_1_, *v* ∈ *V*_1_} and *E*_*t*_ (*V*_1_) = {(*u*, *v*) ∈ *E* | *u* ∈ *V*_1_, *v* ∈ *V* \ *V*_1_}. Following Proposition 6, we have the following result.

#### Proposition 7

*Let* 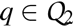. *We have that* 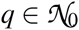 *if and only if there exists a connected subset V*_1_ ⊆ *V*_0_ *such that E_s_*(*q*) = *E*_*s*_(*V*_1_) *and E_t_* (*q*) = *E*_*t*_ (*V*_1_), or *E*_*s*_(*q*) = *E*_*t*_ (*V*_1_) *and E_t_* (*q*) = *E*_*s*_(*V*_1_).

Proposition 7 transforms the problem of deciding whether a vector 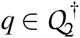 is also in 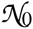 into the problem of examining whether *E*_*s*_(*q*) and *E*_*t*_ (*q*) form a set of cut edges, which can be accomplished through two rounds of breadth-first searching (Algorithm 2).

#### Algorithm 2 Exact algorithm to compute 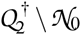

**Input:** 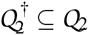, *G*

**Output:** 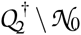

1. **FOREACH** 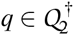, **DO** the following procedure for both *q*′ = *q* and *q*′ = −*q*

2. Traverse *G* from edges in *E*_*s*_ (*q*′), stop searching when reaching edges in *E*_*t*_ (*q*′). **IF** all searching ends up with edges in *E*_*t*_ (*q*′), i.e., *t* is not reached, **THEN** add *q* to 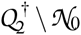.

3. Traverse *G* reversely from edges in *E*_*t*_ (*q*′), stop searching when reaching edges in *E*_*s*_ (*q*′). **IF** all searching ends up with edges in *E*_*s*_ (*q*′), i.e., *s* is not reached, **THEN** add *q* to 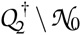.

Algorithm 2 also runs in *O*(|*E* | · | *f* |) time, since 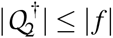 and for each 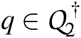, Algorithm 2 takes *O*(|*E*|) time to check its membership in 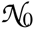. Thus, our heuristic for Problem 2, which consists of Algorithm 1 followed by Algorithm 2, also runs in *O*(|*E* | · | *f* |) time.

### 3.2 Reducing the Optimality Gap

In this section, we devise a heuristic (Algorithm 3) to use 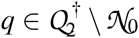 to reduce the optimality gap. The idea of is to iteratively choose one edge from *E*_*s*_(*q*) and one edge from *E*_*t*_ (*q*), merge them through updating *G*, *f* and *q*, until finally q becomes 0.

#### Algorithm 3 Heuristic to reduce the optimality gap

**Input:** 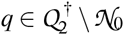, *G* and *f*

**Output: TRUE** and updated *G* and *f* if the optimality gap is reduced, **FALSE** otherwise

1. **WHILE** *q* ≠ 0

2. Arbitrarily choose *e*_*i*_ = (*u*_1_, *v*_1_) ∈ *E*_*s*_ (*q*) and *e*_*j*_ = (*u*_2_, *v*_2_) ∈ *E*_*t*_ (*q*) that are connected; if no such pair can be found, **RETURN FALSE**. Assume a path from *v*_1_ to *u*_2_ exists; also assume *f* (*e*_*i*_) ≤ *f* (*e*_*j*_).

3. **IF** *v*_1_ = *u*_2_ (see Figure 3**(a)**)

4. Add edge *e*_*k*_ = (*u*_1_, *v*_2_) to *G* with flow value of *f* (*e*_*i*_); set *q*[*k*] = 0; remove *e*_*i*_; set *q*[*i*] = 0.

5. Remove *e _j_* and set *q*[ *j*] = 0 if *f* (*e*_*i*_) = *f* (*e _j_*), otherwise set *f* (*e _j_*) as *f* (*e _j_*) − *f* (*e*_*i*_).

6. **ELSE** (see Figure 3**(b)**)

7. Apply Algorithm 6 (Section 3.3) to make *e*_*i*_ and *e _j_* adjacent. **IF** this succeeds, **GOTO** step 3.

8. Arbitrarily choose a path *p* from *v*_1_ to *u*_2_ satisfying that all edges in *p* have a flow value of at least *f* (*e*_*i*_). We assume that *p* does not contain edges in *E*_*s*_ (*q*) ∪ *E*_*t*_ (*q*), otherwise we switch to merge this closer pair of edges. **IF** such path cannot be found, **RETURN FALSE**.

9. Add edge *e*_*k*_ = (*u*_1_, *v*_2_) to *G* with flow value of *f* (*e*_*i*_); set *q*[*k*] = 0; remove *e*_*i*_; set *q*[*i*] = 0.

10. For edge *e _j_* and every edge in *p*, subtract their flow value by *f* (*e*_*i*_); if their updated flow values become 0, then remove them from the graph and set *q*[·] = 0 for them.

11. **RETURN TRUE**.

Suppose that Algorithm 3 succeeds after n iterations. Let *q*^*k*^, *G*^*k*^ = (*V*^*k*^, *E*^*k*^) and *f*^*k*^ be the updated vector, graph, and flow, respectively, after it finishes *k* iterations, 1 ≤ *k* ≤ *n*. Notice that *q*^*n*^ = 0. Similar to 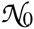 and 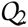, we also define 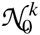 and 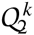 w.r.t. *G*^*k*^ and *f*^*k*^. Let (*P*^*k*∗^, *w*^*k*∗^) be one optimal decomposition of (*G*^*k*^, *f*^*k*^), and also defineΔ^*k*^ = |*E*^*k*^| − |*V*^*k*^ | + 2, 1 ≤ *k* ≤ *n*. Assume that *G*^0^ = *G* and *f* ^0^ = *f* for sake of simplicity.

**Figure 3:**
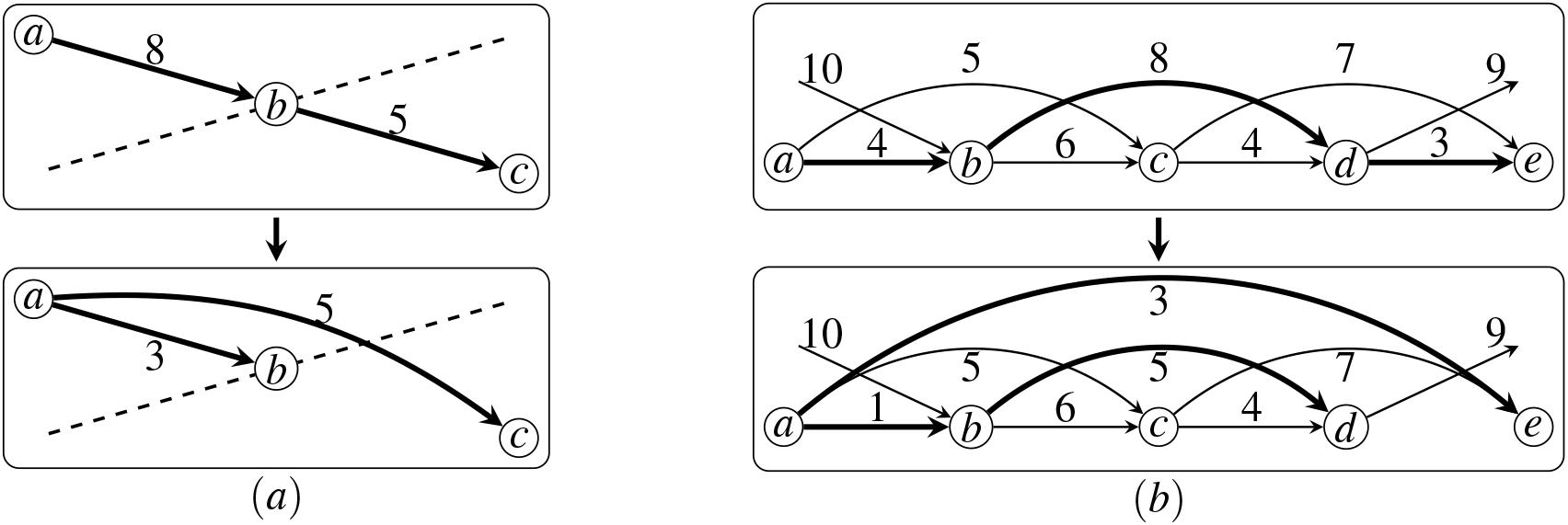
**(a)** Merging two adjacent edges. **(b)** Merging two distant edges (*a*, *b*) and (*d*, *e*).

#### Proposition 8

*If Algorithm 3 succeeds, then we have* 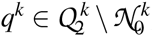, 1 ≤ *k* ≤ *n* − 1.

#### Proposition 9

*If Algorithm 3 succeeds, then we have n* = |*E*_*s*_(*q*)| + |*E*_*t*_ (*q*)| − 1.

We emphasize that the merging process in Algorithm 3 narrows down the solution space. Specifically, the set of all possible decompositions of (*G*^*k*^, *f*^*k*^) is a subset of that of (*G*^*k*−1^, *f* ^*k*−1^), 1 ≤ *k* ≤ n − 1, i.e., every decomposition of (*G*^*k*^, *f ^k^*) corresponds to a decomposition of (*G*^*k*−1^, *f*^*k*−1^). This implies that |*P*^*k*^ ^∗^| ≥ |*P*^(*k*−1)^ ^∗^|. Notice that it is possible that |*P*^*k*∗^| > |*P*^(*k*−1)∗^|, i.e., optimal solutions get eliminated during Algorithm 3. Figure 4 shows that merging two adjacent edges (lines 3–5 of Algorithm 3) might break optimality.

**Figure 4:**
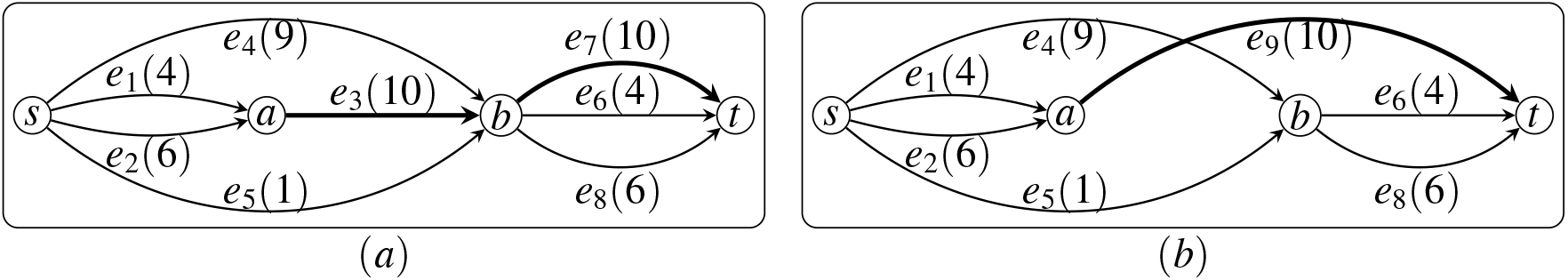
Merging adjacent edges might break optimality. **(a)** Original graph with 4 paths in the optimal decomposition. **(b)** Graph after merging *e*_3_ and *e*_7_, with 5 paths in the optimal decomposition.

We now describe our central result, which states that if Algorithm 3 succeeds, then the optimality gap will be reduced by at least 1. The intuition behind the proof is that during the first (*n* − 1) iterations, (|*E*| − |*V*|) keeps unchanged or decreases, while at the last iteration, this value shall be reduced by at least 1.

#### Proposition 10

*If Algorithm 3 succeeds, then we have* Δ^*n*^ − |*P*^*n*∗^| ≤ − |*P*^∗^| − 1.

### 3.3 Making Two Distant Edges Adjacent

We now describe an algorithm (Algorithm 6) to try to make two given distant edges adjacent while keeping optimality, a key subroutine used in Algorithm 3 (line 7). In this Section, we describe the intuition and idea of this algorithm, and its formal and complete description is in Appendix A. We first characterize the hierarchical structure of the graph. We say a pair of vertices *u* and *v* is *closed*, if there does not exist any edge satisfying that exactly one of its ends is in the subgraph induced by these two vertices (see Figure 5 **(a,c)**).

**Figure 5:**
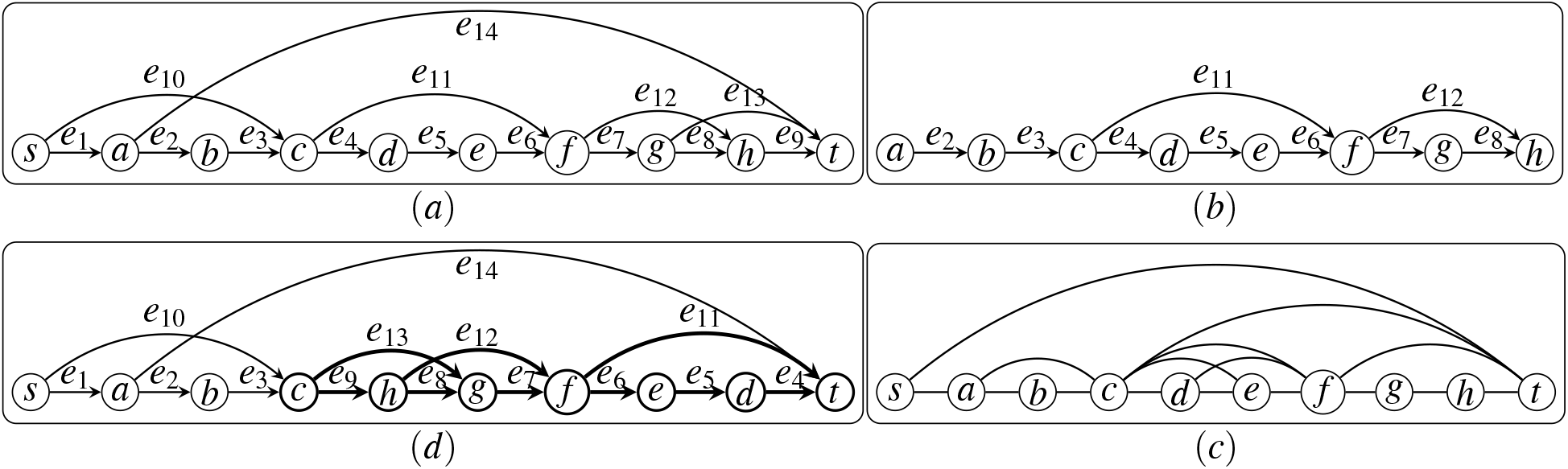
**(a)** Illustration of *G* = (*V*, *E*). **(b)** Illustration of *V* (*a*, *h*) and *E* (*a*, *h*). **(c)** Edges represent all closed pairs (i.e., 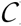). **(d)** Graph after reversing closed pair of (*c*,*t*). Notice that edges {*e*_3_, *e*_10_} and {*e*_9_, *e*_13_} become adjacent, which are distant before reversing.

With a given pair of closed vertices, we then define a *reverse* operation to modify the graph (Figure 5 **(a,d)**). We can prove that, performing reverse operations will not change the optimality (Proposition 12). Thus, we focus on the problem of deciding whether there exists a series of reverse operations to make two given distant edges adjacent (Problem 3).

We give a polynomail-time exact algorithm to solve Problem 3 (Algorithm 6). The algorithm first use a DAG to organize all closed pairs (Figure 6). Then through traversing the DAG starting from the vertices corresponding to the two given distant edges, we can decide whether these two edges can be made adjacent, and if so the reverse operations can be retrieved by tracing back the DAG.

**Figure 6:**
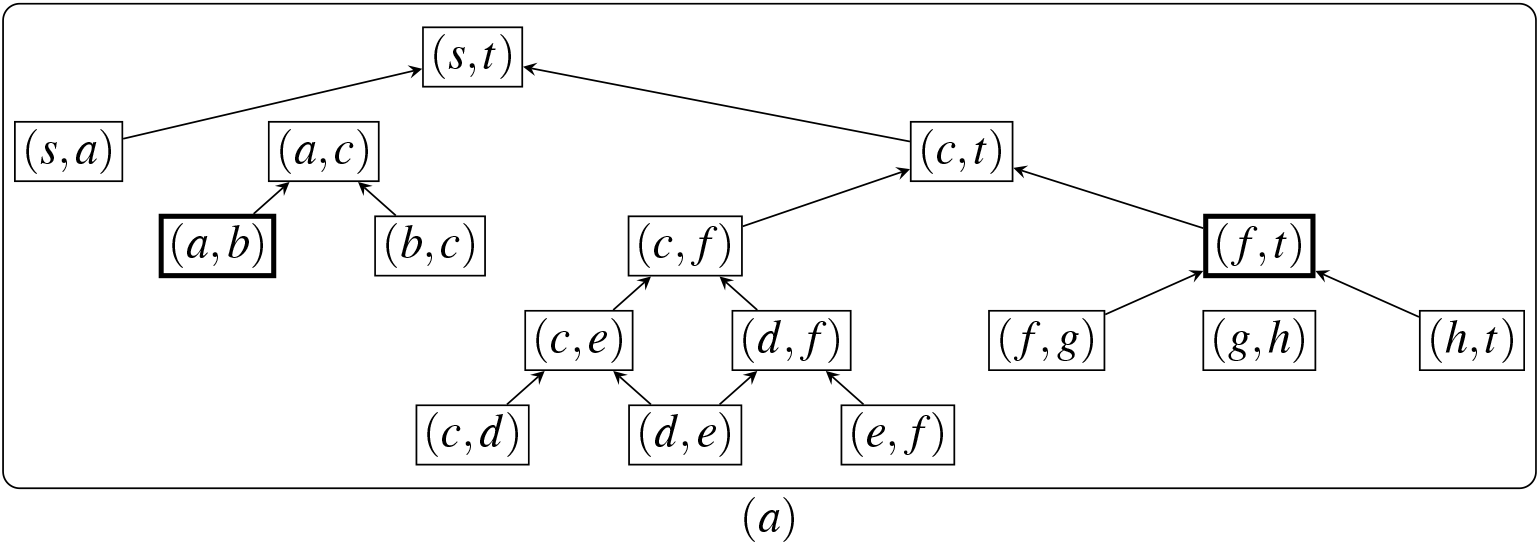
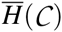 w.r.t. the graph in Figure 5 **(a)**. To make edge *e*_*i*_ = (*a*, *b*) and edge *e*_*j*_ = (*f*, *h*) adjacent, Algorithm 6 first computes their closures in 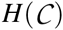, which are (*a*, *b*) and (*f*,*t*) respectively. The following traversing procedure gives *S*_1_ = {*a*}, *T*_1_ = {*b*, *c*}, *S*_2_ = { *f*, *c*, *s*}, and *T*_2_ = {*t*}. We have *T*_1_ ∩ *S*_2_ = *c*, i.e., *e*_*i*_ and *e*_*j*_ can be made adjacent at *c*, and the operations are reversing (*a*, *c*), (*f*,*t*) and (*c*,*t*).

### 3.4 The Complete Algorithm

With all above components available, we now formally state our heuristic for Problem 1 in Algorithm 4. The first phase (lines 1 to 3) is to iteratively reduce the optimality gap. It then uses the greedy-width algorithm (proposed in [24]) to fully decompose the remaining graph into paths. Notice that these paths are *w.r.t*. the updated graph, and we need to recover the paths corresponding to the original graph (we maintain the tracing back information when updating the graph).

#### Algorithm 4 Heuristic for Problem 1

**Input:** *G* and *f*

**Output:** A decomposition (*P*^†^, *w*^†^) of (*G*, *f*)

1. Apply Algorithm 1 and Algorithm 2 to compute 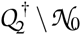. **IF** 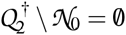, **GOTO** step 4.

2. **FOREACH** 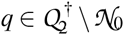

3. Apply Algorithm 3 with *q*. **IF** it succeeds, **GOTO** step 1.

4. Apply the greedy-width algorithm on the updated graph.

5. For the paths returned in line 4, recover the corresponding paths *w.r.t*. the original graph.

#### Proposition 11

*Algorithm 4 runs in O*(|*V* |^2^ · |*E* |^2^ · | *f* |) *time*.

From [24], we know that for an instance with optimality gap of 0, the greedy-width algorithm can guarantee returning one optimal decomposition. This implies that, for any instance, if the first phase of Algorithm 4 closes its optimality gap and does not break the optimality, then Algorithm 4 shall return an optimal decomposition for this instance. We use experimental studies to illustrate its performance in Section 4.

## 4 Experimental Results

We first describe the experiments in the context of transcript assembly. We use four datasets. The first dataset includes 1000 human RNA-seq samples from SRA (Sequence Reads Archive). For each gene in each sample, we build an instance (i.e., *G* and *f*) as follows. We first use Salmon [30] to identify and quantify the expressed transcripts of this gene: the expressed transcripts shall be then used as the ground truth paths, denoted as *P*^‡^, while the expression abundances of these transcripts shall be used as the ground truth weights, denoted as *w*^‡^. With (*P*^‡^, *w*^‡^), we then construct *G* and *f*: vertices of *G* correspond exons, edges are the union of the edges in the ground truth paths, and the flow value for each edge is the superposition of the ground truth weights of the paths that go through this edge (Figure 7). For the generated instance (*G*, *f*), we pipe it into the two algorithms (Algorithm 4 and greedy-width). We denote by (*P*^†^, *w*^†^) as the predicted paths and weights by any of these two algorithms. We say (*P*^†^, *w*^†^) is *correct* if we have |*P*^†^| ≤ |*P*^‡^|. Notice that (*P*^‡^, *w*^‡^) is not necessarily optimal, but only gives an upper bound, i.e., |*P*^‡^| ≥ |*P*^∗^|. The other three datasets are obtained by simulation using Flux-Simulator [31] on three well-annotated species, human, mouse and zebrafish. For each species, we independently simulate 100 samples. For each gene in each sample, we collect the ground truth (*P*^‡^, *w*^‡^) from Flux-Simulator, and use them to build the instance *G* and *f* (Figure 7).

**Figure 7:**
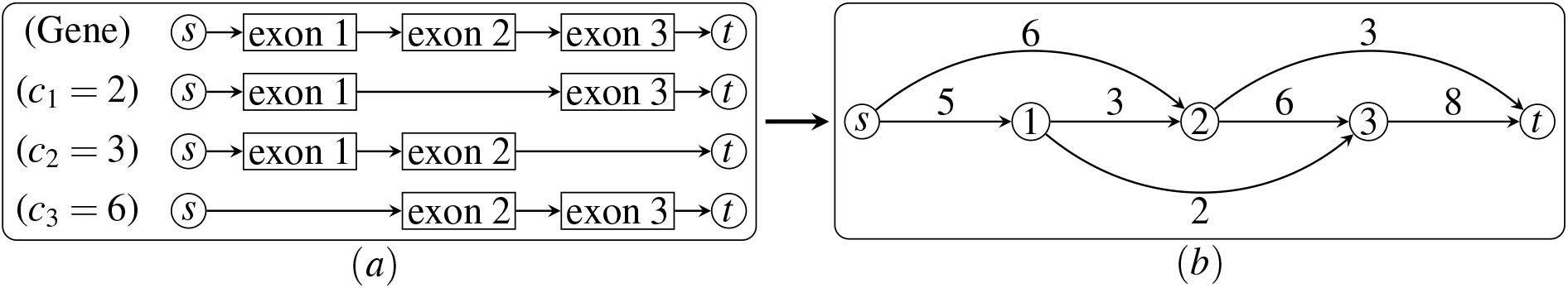
**(a)** Transcripts and their expression abundances (*c*_*k*_) of a gene. **(b)** The splice graph *G* and flow *f* derived from the ground truth in **(a)**.

Based on |*P*^‡^|, for each dataset, we classify all instances into 10 categories, i.e., |*P*^‡^| ∈ {1, 2, …, 10}, and remove those instances with |*P*^‡^| > 10 (which is a tiny portion). For each category, we run both algorithms on all instances and compare their *accuracy* (proportion of instances that are predicted correctly), and average ratio between |*P*^†^| and |*P*^‡^| (i.e., mean value of |*P*^†^|/|*P*^‡^| over all instances).

The results are shown in Figure 8. We can observe that Algorithm 4 gives very high accuracy on all instances, while the accuracy of the greedy-width algorithm decreases rapidly as |*P*^‡^| increases. Meanwhile, the decompositions given by the greedy-width algorithm uses more paths and the ratio between the number of predicted paths and the ground truth keeps increasing as |*P*^‡^| increases, while for Algorithm 4 such ratio constantly remains closing to 1. These results demonstrate that in the application of transcript assembly, Algorithm 4 significantly outperforms the greedy-width algorithm.

**Figure 8:**
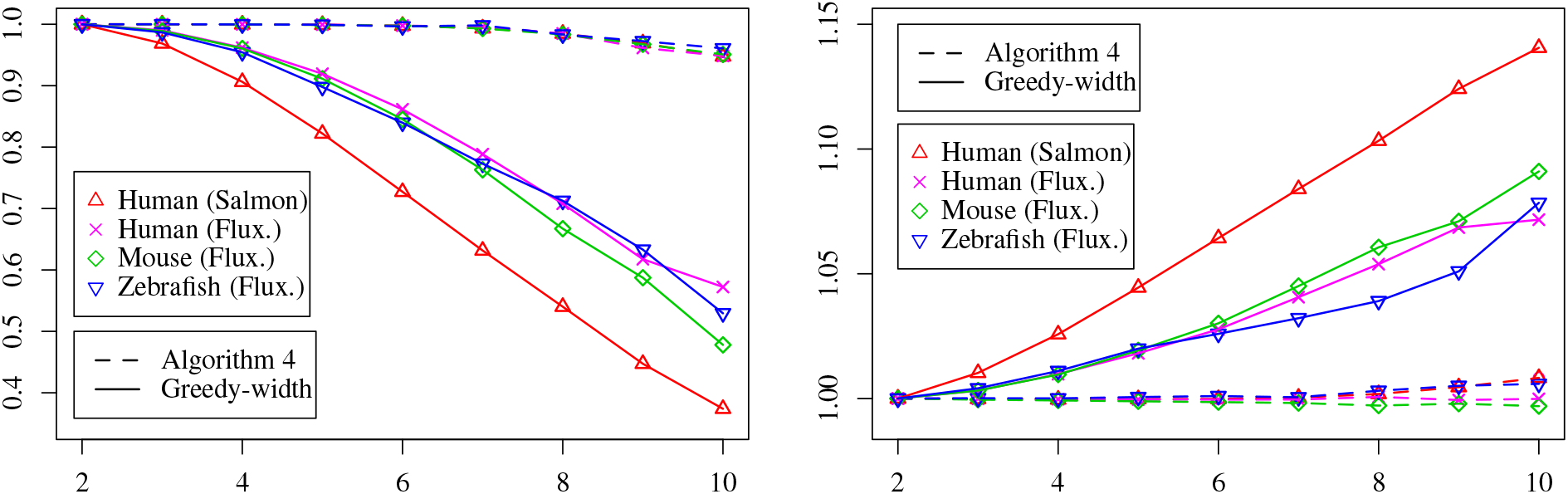
Comparison of the two algorithm on transcript assembly. **Left:** *x*-axis is |*P*^‡^| and *y*-axis is the accuracy. **Right:** *x*-axis is |*P*^‡^| and *y*-axis is the average ratio between |*P*^†^|/|*P*^‡^|.

To evaluate the performance of our algorithm on large graphs, we carry out another set of experiments using simulated random instances. The results show that our algorithm can give very high accuracy for a wide spectrum of parameter combinations, while the greedy-width algorithm works well only for easy instances. These results further demonstrate the significant improvement of our algorithm over the greedy-width algorithm. The simulation setup and the detailed results are in Appendix B.

## 5 Discussion

We give an efficient heuristic for the minimum path flow decomposition problem. The components of this heuristic can be further improved. For example, we shall study how to reduce the optimality gap while keep optimality if a true nontrivial null vector is given, and also study how to identify nontrivial null vectors that are not simple as well as how to use such vectors to reduce the optimality gap. We tackle here only the flow decomposition step of transcript assebmly. Future work also includes extending our algorithm to develop a complete transcript assembler.

## Acknowledgements

We thank Meiyue Shao and Guillaume Marc¸ais for helpful discussions and suggestions. This research is funded in part by the Gordon and Betty Moore Foundation’s Data-Driven Discovery Initiative through Grant GBMF4554 to Carl Kingsford, by the US National Science Foundation (CCF-1256087, CCF-1319998) and by the US National Institutes of Health (R21HG006913, R01HG007104). C.K. received support as an Alfred P. Sloan Research Fellow.

## Appendix A Making Two Distant Edges Adjacent

In this Section, we describe an algorithm to make two given distant edges adjacent while keeping optimality. We first give some definitions to characterize the hierarchical structure of *G*. We say a pair of vertices (*u*, *v*) is *connected* (*w.r.t.* graph *G*), if there exists a path from *u* to *v* in *G*. For a pair of connected vertices (*u*, *v*), we define a subset *V* (*u*, *v*) ⊆ *V* as the set of vertices that are on some path from u to v (see Figure 5 **(a,c)**). Notice that if (*u*, *v*) is connected, we always have {*u*, *v*} ⊆ *V* (*u*, *v*). For a pair of connected vertices (*u*, *v*), we also define a subset *E* (*u*, *v*) ⊆ *E* as follows (see Figure 5): for any *e* ∈ *E* we say *e* ∈ *E* (*u*, *v*) if the both ends of *e* are in *V* (*u*, *v*). We say a pair of connected vertices (*u*, *v*) is *closed*, if for every vertex *w* ∈ *V* (*u*, *v*) \ {*u*, *v*}, all incident edges of *w* in *E* are also in E (*u*, *v*). We denote by 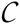 the set of all closed pairs. For each pair 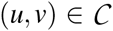, we define *E*_*s*_(*u*, *v*) ⊆ *E* (*u*, *v*) as the set of edges in *E* (*u*, *v*) that are adjacent to *u*, and define *E*_*t*_ (*u*, *v*) ⊆ *E* (*u*, *v*) as the set of edges in *E* (*u*, *v*) that are adjacent to *v*.

We define a *reverse* operation to modify *G* (Figure 5 **(a,d)**). For each pair 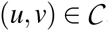, we can reverse it by first detaching edges in *E*_*s*_(*u*, *v*) from u and attaching them to *v*, detaching edges in *E*_*t*_ (*u*, *v*) from *v* and attaching them to *u*, and then switching the direction of all edges in *E* (*u*, *v*). Notice that we treat the reverse operation in a way that the set of edges in *G* is kept unchanged, but the locations and orientations of edges might be changed. Formally, let *G*′ be the graph after reversing 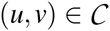. We assume that *G*′ still have the edge set of *E*. But for each edge *e*_*i*_ = (*u*, *v*) in *G*, it might become *e*_*i*_ = (*u*′, *v*′) in *G*′ with *u*′ ≠ *u* and/or *v*′ ≠ *v*. In other words, all edges are recognized by their original identities in *E*, and their adjacent two vertices might become different after performing reverse operations.

Let *G*′ be the graph after reversing 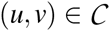. Let (*P*′^∗^, *w*′^∗^) be one optimal decomposition of (*G*′, *f*). The following result states that solving Problem 1 on (*G*, *f*) is equivalent to solving it on (*G*′, *f*).

### Proposition 12

*There exists a one-to-one correspondence between all decompositions of* (*G*, *f*) *and all decompositions of* (*G*′, *f*). *In particular, we have that* |*P*^∗^| = |*P*′^∗^|.

Based on Proposition 12, we address the following problem.

### Problem 3

*Given two edges e_i_*, *e*_*j*_ ∈ *E*, *decide whether we can perform a series of reverse operations, such that in the resulting graph we have e_i_* = (*u*_1_, *v*_1_) and *e _j_* = (*u*_2_, *v*_2_) *with either v*_1_ = *u*_2_ or *v*_2_ = *u*_1_.

To solve Problem 3, we first describe a data structure to organize all the reverse operations. We define a binary relation 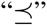 on 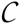: for two pairs (*u*_1_, *v*_1_), 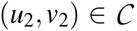, we say 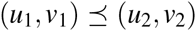 if *V* (*u*_1_, *v*_1_) ⊆ *V* (*u*_2_, *v*_2_). It is easy to verify that this relation forms a partial order. Let 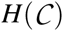 be the Hasse diagram on 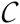 *w.r.t*. this partial order. For each edge in 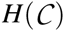 from (*u*_1_, *v*_1_) to (*u*_2_, *v*_2_), we remove it from 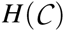 if *u*_1_ ≠ *u*_2_ and *v*_1_ ≠ *v*_2_. We denote by 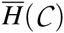 this updated graph (see Figure 6).

We now describe an exact algorithm to solve Problem 3, which requires constructing 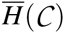 and then search-ing series of reverse operations guided by 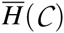. We first compute the transitive closure of *G*, which gives us all possible connected pairs. For each connected pair (*u*, *v*), we follow the definition to check whether this pair is in 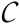: we compute *V* (*u*, *v*), which can be done in *O*(|*V* |) time with the availability of all connected pairs; then for each vertex *w* ∈ *V* (*u*, *v*) \ {*u*, *v*} we check whether all adjacent edges of w are still in *V* (*u*, *v*). It takes *O*(|*E* |) time to verify each connected pair, and thus the total running time to compute 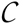 is *O*(|*V* |^2^ · |*E* |).

We then give an algorithm to compute 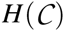, i.e., for each pair in 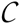 we need to compute its parents node in 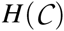. For a subset *V*_1_ ⊆ *V*, |*V*_1_| ≥ 2, we say 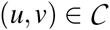 is a closure of *V*_1_, if we have *V*_1_ ⊆ *V* (*u*, *v*), and there does not exist 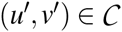 such that *V*_1_ ⊆ *V* (*u*′, *v*′) ⊂ *V* (*u*, *v*). Intuitively, the closure of a given set of vertices is the smallest closed pair whose set of vertices contains the given vertices. As we will see in Proposition 13 such closure is unique. Now we describe an exact algorithm to compute the closure of a given subset, which shall be used in both constructing 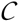 and the following steps (line 2 of Algorithm 6). The idea of this algorithm is to iteratively add necessary vertices until finally it becomes closed.

### Algorithm 5 Exact algorithm to compute closure

**Input:** *V*_1_ ⊂ *V*, 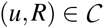 and *G*

**Output:** Closure of *V*_1_

1. Compute a topological ordering *O* of *V*, i.e., if (*u*, *v*) is connected, then *O*(*u*) < *O*(*v*).

2. Let *S* be the set containing all vertices in the final closure, initialized as *V*_1_.

3. Let *L* and *R* be the smallest and largest vertex in *S* according to their orderings in *O*, respectively.

4. Let *Q* be a queue that stores all (unexamined) vertices in *S* \ {*L*, *R*}, initialized as *S* \ {*L*, *R*}.

5. **REPEAT**

6. **WHILE** *Q* is not empty

7. Pop element v from *Q*.

8. **FOREACH** adjacent vertex *u* of *v* that is not in *S*

9. Insert *u* into *S*.

10. **IF** *O*(*u*) < *O*(*L*), push *L* to *Q* and set *L* = *u*.

11. **IF** *O*(*u*) > *O*(*R*), push *R* to *Q* and set *R* = *u*.

12. **FOREACH** adjacent vertex *u* of *L* that is not in *S*

13. **IF** 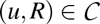, insert *u* into *S* and push *u* to *Q*.

14. **IF** *Q* is empty, **RETURN** (*L*, *R*).

### Proposition 13

*For any V*_1_ ⊂ *V*, |*V*_1_| ≥ 2, *Algorithm 5 returns the unique closure of V*_1_ *in O*(|*E* |) *time*.

We now describe how to build 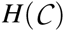 using Algorithm 5. Suppose that (*u*′, *v*′) is one of the direct parents of 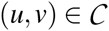 in 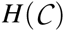. We have two cases: either *u* ∉ {*u*′, *v*′}, or *v* ∉ {*u*′, *v*′}. If it is the first case, then (*u*′, *v*′) is exactly the closure of *V*_1_ = {*u*, *v*} ∪ {*w* | (*w*, *u*) ∈ *E*} ∪ {*w* | (*u*, *w*) ∈ *E*}. If it is the second case, then (*u*′, *v*′) is exactly the closure of *V*_2_ = {*u*, *v*} ∪ {*w* | (*w*, *v*) ∈ *E*} ∪ {*w* | (*v*, *w*) ∈ *E*}. In other words, the parents of 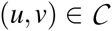 in 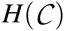 are exactly the closures of *V*_1_ and *V*_2_. For each pair 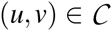, we use this procedure to compute its parents, and thus the whole graph 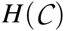 can be constructed. To further build 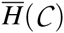, by definition, we then check each edge of 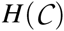 to see whether the two pairs corresponding to its two adjacent vertices share the same starting or ending vertex. The total running time to build 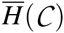 is *O*(|*V* |^2^ · |*E* |).

### Algorithm 6 Exact algorithm for Problem 3

**Input:** *G*, *e*_*i*_ = (*u*_1_, *v*_1_), *e*_*j*_ = (*u*_2_, *v*_2_) ∈ *E*

**Output: TRUE** and series of reverse operations if they can be made adjacent, **FALSE** otherwise

1. Compute 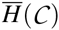.

2. Compute the closures of {*u*_1_, *v*_1_} and {*u*_2_, *v*_2_}; denote them as (*u*′_1_, *v*′_1_) and (*u*′_2_, *v*′_2_), respectively.

3. Initialize *S*_1_ = {*u*_1_}, *T*_1_ = {*v*_1_}, *S*_2_ = {*u*_2_}, *T*_2_ = {*v*_2_}.

4. Traverse 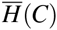 from (*u*′_1_, *v*′_1_): for each visited node 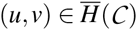, insert *u* to *S*_1_ and insert *v* to *T*_1_.

5. Traverse 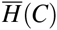 from (*u*′_2_, *v*′_2_): for each visited node 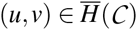, insert *u* to *S*_2_ and insert *v* to *T*_2_.

6. **IF** *S*_1_ ∩ *T*_2_ = ∅ and *S*_2_ ∩ *T*_1_ = ∅, **RETURN FALSE**.

7. Compute arbitrarily *w* ∈ (*S*_1_ ∩ *T*_2_) ∪ (*S*_2_ ∩ *T*_1_); assume that *w* ∈ *S*_2_ ∩ *T*_1_.

8. Compute any path from (*u*′_1_, *v*′_1_) to node (·, *w*), and traverse this path to construct the reverse operations to make *e*_*i*_ point to *w*: let (*u*, *v*) be the current node (i.e., *e*_*i*_ is either points to *v* or leaves *u*) and *u* be the boundary vertex shared by (*u*, *v*) and its parent; reverse (*u*, *v*) if *e*_*i*_ is not adjacent to *u*; move to the parent of (*u*, *v*) on this path. In the end reverse (·, *w*) if *e*_*i*_ is not adjacent to *w*.

9. Perform the same procedure as line 8 to construct the reverse operations to make *e*_*j*_ leave *w*.

10. **RETURN TRUE**.

Our complete algorithm to solve Problem 3 is described in Algorithm 6. The idea is first to locate the nodes in 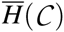 for the given two edges, and then traverse 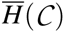 starting from these two nodes respectively to compute all the possible positions that the two given edges can be moved to by reverse operations. After that the algorithm examines whether there exists a common vertex where the two edges can be made adjacent. The series of reverse operations can be obtained through tracing back. The correctness and running time of Algorithm 6 is stated as Proposition 14.

### Proposition 14

*Algorithm 6 gives an exact solution for Problem 3 in O*(|*V* |^2^ · |*E* |) *time*.

## Appendix B Experiments with Simulated Random Instances

The simulation procedure first generates the ground truth (*P*^‡^, *w*^‡^), and then builds (*G*, *f*) based on them. In the simulation, we assume that *G* = (*V*, *E*) contains 1000 vertices, i.e., |*V* | = 1000, and assume that (1, 2, …, |*V* |) is one topological sorting of *G*. To generate a path *p*^‡^ in the ground truth, we first fix the length of this path, which is sampled uniformly at random from {1, 2, …, *L*}, where *L* ≤ |*V* | − 1 is a parameter governing the maximum length of the paths in the ground truth path. We then sample (|*p*^‡^| + 1) vertices uniformly at random without replacement from {1, 2, …, |*V* |}, and the path connects these vertices in an ascending order. The ground truth weight of this path, i.e., *w*^‡^(*p*^‡^), is sampled uniformly at random from {1, 2, …, 10000}. We repeat this process to generate |*P*^‡^| paths, where |*P*^‡^| is another parameter governing the number of paths in the ground truth. With (*P*^‡^, *w*^‡^), we then construct instance (*G*, *f*) by superposition of the weights of the paths that go through the edge (similar to Figure 7). Again, we denote by (*P*^†^, *w*^†^) be the output of the algorithm (Algorithm 4 or greedy-width), and we say (*P*^†^, *w*^†^) is *correct* if we have |*P*^†^| ≤ |*P*^‡^|.

We perform experiments with two different parameter settings, one is *L* = 50, |*P*^‡^| ∈ {20, 40, …, 200}, the other is *L* ∈ {10, 20, …, 100}, |*P*^‡^| = 100. For each parameter combination (*L*, |*P*^‡^|), we randomly generate 100 instances using the procedure mentioned above. We run both algorithms on these 100 instances, and for each algorithm we record its *accuracy* (proportion of instances that are predicted correctly), average ratio between |*P*^†^| and |*P*^‡^| (i.e., mean value of |*P*^†^|/|*P*^‡^| over the 100 instances), and total running time.

The results for the first parameter settings are shown in Table 1. First, we can observe that Algorithm 4 gives 100% accuracy when |*P*^‡^| ≤ 140, and it decreases a few percentage for larger |*P*^‡^|. Meanwhile, the accuracy of the greedy-width algorithm decreases rapidly with the increase of |*P*^‡^|, and drops to 0 when |*P*^‡^| ≥ 100. Second, we can see that the number of paths returned by Algorithm 4 is very close to the ground truth for all instances, while greedy-width algorithm uses more paths than the ground truth especially for large |*P*^‡^|. Third, both algorithms can finish these instances in a short amount of time, and Algorithm 4 is about 4 times slower than the greedy-width algorithm.

**Table 1:** Comparison of the two algorithms on simulations with *L* = 50.

| | $\vert P^\ddagger\vert $ | 20 | 40 | 60 | 80 | 100 | 120 | 140 | 160 | 180 | 200 |
| --- | --- | --- | --- | --- | --- | --- | --- | --- | --- | --- | --- |
| Accuracy (%) | Algorithm 4 | 100 | 100 | 100 | 100 | 100 | 100 | 100 | 97 | 93 | 83 |
|  | Greedy-width | 94 | 46 | 14 | 1 | 0 | 0 | 0 | 0 | 0 | 0 |
| Mean $\vert P^\dagger\vert /\vert P^\ddagger\vert $ | Algorithm 4 | 1.00 | 1.00 | 1.00 | 1.00 | 1.00 | 1.00 | 1.00 | 1.00 | 1.00 | 1.01 |
|  | Greedy-width | 1.01 | 1.01 | 1.10 | 1.21 | 1.27 | 1.40 | 1.51 | 1.66 | 1.76 | 1.83 |
| Time (seconds) | Algorithm 4 | 4 | 14 | 30 | 51 | 72 | 91 | 146 | 193 | 223 | 255 |
|  | Greedy-width | 2 | 5 | 8 | 13 | 17 | 23 | 33 | 44 | 55 | 65 |

The results for the second parameter settings are shown in Table 2, which are consistent with that in Table 1. Combining these results we can conclude that Algorithm 4 can give decompositions with the same number of paths as that in the ground truth for a large scope of parameter combinations, while greedy-width algorithm can only give such decompositions for small *L* and small |*P*^‡^|, and it requires a significant portion of more paths for large *L* or large |*P*^‡^|. This shows a considerable improvement of Algorithm 4 over the greedy-width algorithm, especially for instances with many paths.

**Table 2:** Comparison of the two algorithms on simulations with |*P*^‡^| = 100.

| $L$ | | 10 | 20 | 30 | 40 | 50 | 60 | 70 | 80 | 90 | 100 |
| --- | --- | --- | --- | --- | --- | --- | --- | --- | --- | --- | --- |
| Accuracy (%) | Algorithm 4 | 100 | 100 | 100 | 100 | 100 | 100 | 100 | 99 | 91 | 81 |
|  | Greedy-width | 65 | 20 | 3 | 0 | 0 | 0 | 0 | 0 | 0 | 0 |
| Mean $\vert P^\dagger\vert /\vert P^\ddagger\vert $ | Algorithm 4 | 1.00 | 1.00 | 1.00 | 1.00 | 1.00 | 1.00 | 1.00 | 1.00 | 1.00 | 1.01 |
|  | Greedy-width | 1.01 | 1.02 | 1.07 | 1.17 | 1.29 | 1.45 | 1.61 | 1.82 | 2.05 | 2.25 |
| Time (seconds) | Algorithm 4 | 5 | 16 | 32 | 54 | 67 | 89 | 146 | 167 | 195 | 226 |
|  | Greedy-width | 3 | 6 | 9 | 13 | 17 | 22 | 31 | 41 | 49 | 58 |

## Appendix C Proofs of Propositions

### Proof of Proposition 1.

Because (*P*^∗^, *w*^∗^) forms a decomposition of (*G*, *f*), we know that *f* = *w*^∗^ · *P*^∗^. By definition, for any 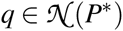 we also have *P*^∗^ · *q* = 0. Thus, we have that *f* · *q* = *w*^∗^ · *P*^∗^ · *q* = 0.

### Proof of Proposition 2.

We can write Pr(*f* · *q* = 0) = Pr(*w*^∗^ · *P*^∗^ · *q* = 0). Since 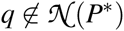, we know that *P*^∗^ · *q* ≠ 0. Combining the fact that elements in *w*^∗^ are sampled independently and uniformly at random from ℤ_*k*_, using the standard backward analysis, we can conclude that Pr(*f* · *q* = 0) ≤ 1/*k*.

### Proof of Proposition 3.

We can easily verify that for every row *p*_*i*_ of 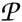 and for every *v* ∈ *V*_0_ we have that *p*_*i*_ · *q*_*v*_ = 0. Particularly, since 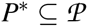, we have that *P*^∗^ · *q*_*v*_ = 0. This implies that 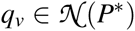, for every *v* ∈ *V*_0_. Hence, we have 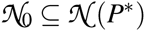.

### Proof of Proposition 4.

According to the rank-nullity theorem, we have that 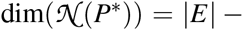 *rank*(*P*^∗^). From [24], we have that rows in the optimal solution are linearly independent, i.e., *rank*(*P*^∗^) = |*P*^∗^|. Besides, we also have that 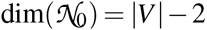, since all vectors in {*q*_*v*_ | *v* ∈ *V*_0_} are linearly independent. Combining all these facts, we have that 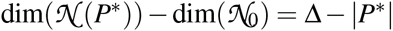.

### Proof of Proposition 5.

We first prove that Algorithm 1 returns a subset of 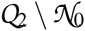. Let *E*_1_ ⊆ *E* and *E*_2_ ⊆ *E* be the two balanced subsets through tracing back from *x* ∈ *D*. We only need to show that *E*_1_ and *E*_2_ are indivisible. We first prove that *E*_1_ and *E*_2_ are disjoint. Suppose conversely that *e* ∈ *E*_1_ ∩ *E*_2_. Then we have *B*[|*E* |, *x* − *f* (*e*)] = 1, since there exist two subsets, namely *E*_1_′ = *E*_1_ \ {*e*} and *E*_2_′ = *E*_2_ \ {*e*}, such that f (*E*_1_′) = *f* (*E*_2_′) = *x* − *f* (*e*), a contradiction to the fact that *x* ∈ *D*.

Now we show that *E*_1_ and *E*_2_ are indivisible. Suppose conversely that there exist strict subsets *E*_1_′ ⊂ *E*_1_ and *E*_2_′ ⊂ *E*_2_ such that *f* (*E*_1_′) = *f* (*E*_2_′) = *y*. Let *e*_*k*_ be the edge with the highest index in *E*_1_ ∪ *E*_2_. Without loss of generality, we assume that *e*_*k*_ ∈ *E*_1_ \ *E*_1_^′^. Let *e*_*k*_′ be the edge with the highest index in *E*_1_′ ∪ *E*_2_′. We have that *B*[*k*′, *y*] = 1, because of the existence of *E*_1_′ and *E*_2_′. Following the recursion of *B* (line 4 of Algorithm 1), we also have that *B*[ *j*, *y* + ∑_*k*_′_<*i*≤ *j*_ *f* (*e*_*i*_)] = 1, for all *k*′ < *j* ≤ *k*. This implies that *B*[*k*, *x* − *f* (*e*_*k*_)] = 1, a contradiction to the fact that *x* ∈ *D*.

We can verify that computing *A*, *B*, *D* and tracing back for all elements from *D* takes *O*(|*E* | · | *f* |) time, while other steps take less than that. Thus, the total running time of Algorithm 1 is *O*(|*E* | · | *f* |).

### Proof of Proposition 6.

On one side, if there exists *V*_1_ ⊆ *V*_0_ such that 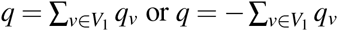, then by definition, we have that 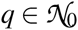.

Now we prove the other side. Assume that 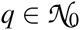, i.e., 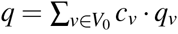. Let *V*_1_ = {*v* ∈ *V*_0_ | *c*_*v*_ ≠ 0}. Let *G*_1_ = (*V*_1_, *E*_1_) be the subgraph of *G* induced by *V*_1_, i.e., we define *e* ∈ *E*_1_ if both ends of e is in *V*_1_. We first prove that *G*_1_ is weakly connected. Suppose conversely that *G*_1_ contains *k* ≥ 2 connected components, 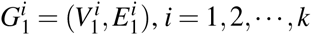. We can write *q* = ∑_1≤*i*≤*k*_ *q*^*i*^, where 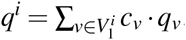. Notice that *q*^*i*^ contributes to distinct elements of *q*, i.e., for any edge 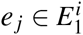, we have *q*[ *j*] = *q*^*i*^[ *j*]. Since 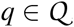, we also have that 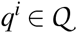. Further, we have that 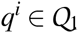, since we have 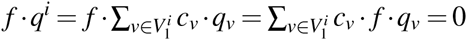. This implies that the two balanced subsets corresponding to *q* can be split into *k* pairs of balanced subsets, namely, *E*_*s*_(*q*^*i*^) and *E*_*t*_ (*q*^*i*^), 1 ≤ *i* ≤ *k*, a contradiction to the fact that 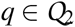.

We now show that *c*_*v*_ ∈ ℤ, ∀*v* ∈ *V*_1_. Let *E*_*s*_(*V*_1_) = {(*u*, *v*) ∈ *E* | *u* ∈ *V* \*V*_1_, *v* ∈ *V*_1_}, and *E*_*t*_ (*V*_1_) = {(*u*, *v*) ∈ *E* | *u* ∈ *V*_1_, *v* ∈ *V* \*V*_1_}, Let *E*_*st*_ (*V*_1_) = *E*_*s*_(*V*_1_) ∪ *E*_*t*_ (*V*_1_). Let *V*_2_ ⊆ *V*_1_ be the subset of vertices that are adjacent to some edge in *E*_*st*_. For any edge *e*_*j*_ = (*u*, *v*) ∈ *E*_1_, we have *q*[ *j*] = *c*_*v*_ − *c*_*u*_. Following this formular, we have that *c*_*v*_ ∈ {−1, +1}, ∀*v* ∈ *V*_2_, since edges in *E*_*st*_ are adjacent to only one vertex in *V*_1_, which is in *V*_2_. Since *G*′ is weakly connected and *q*[ *j*] ∈ {−1, 0, +1}, by propagating from *V*_2_ with the above formular, we can conclude that *c*_*v*_ ∈ ℤ for all *v* ∈ *V*_1_.

Consider any edge *e*_*j*_ = (*u*, *v*) ∈ *E*_1_. If we assume that *c*_*u*_ ≥ 1, then from the above formular we have that *c*_*v*_ = *c*_*u*_ + *q*[ *j*] ≥ 0; and since *c*_*v*_ ≠ 0 and *c*_*v*_ ∈ ℤ, we have that *c*_*v*_ ≥ 1. Following this reasoning and the fact that *G*′ is weakly connected, through propagating we know that {*c*_*v*_ | *v* ∈ *V*_1_} have the same sign. Without loss of generality, we assume that *c*_*v*_ ≥ 1, ∀*v* ∈ *V*_1_. This implies that *c*_*v*_ = 1, ∀*v* ∈ *V*_2_, and further implies that *q*[ *j*] = 1 for *e*_*j*_ ∈ *E*_*s*_(*V*_1_) and *q*[ *j*] = −1 for *e*_*j*_ ∈ *E*_*t*_ (*V*_1_). Thus, we have that *E*_*s*_(*V*_1_) ⊆ *E*_*s*_(*q*) and *E*_*t*_ (*V*_1_) ⊆ *E*_*t*_ (*q*). According to flow conservation, we have that *f* (*E*_*s*_(*V*_1_)) = *f* (*E*_*t*_ (*V*_1_)). Combining these two facts and the assumption that q is indivisible, we must have that *E*_*s*_(*V*_1_) = *E*_*s*_(*q*) and *E*_*t*_ (*V*_1_) = *E*_*t*_ (*q*). This consequently implies that *V*_1_ = *V*_2_. As we already have showed, *c*_*v*_ = 1, ∀*v* ∈ *V*_2_, we have that 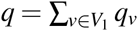.

### Proof of Proposition 7.

Suppose that 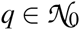. According to Proposition 6, without loss of generality, we have that there exists a connected subset *V*_1_ ⊆ *V*_0_ such that 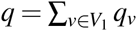. It is clear that *q*[ *j*] = 1 if and only if *e*_*j*_ ∈ *E*_*s*_(*V*_1_), and *q*[ *j*] = −1 if and only if *e*_*j*_ ∈ *E*_*t*_ (*V*_1_), since for any edge *e*_*k*_ that are adjacent to two vertices in *V*_1_ we have *q*[*k*] = 0. This yields that *E*_*s*_(*q*) = *E*_*s*_(*V*_1_) and *E*_*t*_ (*q*) = *E*_*t*_ (*V*_1_).

On the other side, suppose that there exists a subset *V*_1_ ⊆ *V*_0_ such that *E*_*s*_(*q*) = *E*_*s*_(*V*_1_) and *E*_*t*_ (*q*) = *E*_*t*_ (*V*_1_). We now prove that 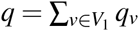. In fact, 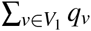 exactly has value of +1 for edges in *E*_*s*_(*V*_1_) = *E*_*s*_(*q*), and has value of −1 for edges in *E*_*t*_ (*V*_1_) = *E*_*t*_ (*q*). This yields that 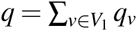.

### Proof of Proposition 8.

According to the algorithm, we can easily verify that 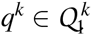, from the fact that 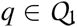. We now show that 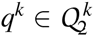. Suppose conversely that *q*^*k*^ is not indivisible, i.e., there exist strict subsets *E*_1_ ⊂ *E*_*s*_(*q*^*k*^) and *E*_2_ ⊂ *E*_*t*_ (*q*^*k*^) such that *f*^*k*^ (*E*_1_) = *f*^*k*^ (*E*_2_). Let *e*_*i*_ ∈ *E*_*s*_(*q*^*k*−1^) and *e*_*j*_ ∈ *E*_*t*_ (*q*^*k*−1^) be the two chosen edges in iteration *k*. Assume that *f*^*k*−1^(*e*_*i*_) ≤ *f* ^*k*−1^(*e*_*j*_). This implies that *e*_*i*_ ∉ *E*_*s*_(*q*^*k*^). Let *E*_1_′ = *E*_1_ ∪ {*e*_*i*_}. If we further have *f* ^*k*−1^(*e*_*i*_) = *f*^*k*−1^(*e*_*j*_), we define *E*_2_′ = *E*_2_ ∪ {*e*_*j*_}, otherwise, we define *E*_2_′ = *E*_2_. Clearly, we have that *f*^*k*−1^(*E*_1_′) = *f* ^*k*−1^(*E*_2_′) according to the algorithm, and that *E*_1_′ and *E*_2_′ are strict subsets of *E*_*s*_(*q*^*k*−1^) and *E*_*t*_ (*q*^*k*−1^), respectively, following from the fact that *E*_1_ and *E*_2_ are strict subsets of *E*_*s*_(*q*^*k*^) and *E*_*t*_ (*q*^*k*^), respectively. Thus, we conclude that *q*^*k*−1^ is not indivisible, a contradiction.

We then show that 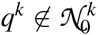. Suppose conversely that 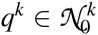. According to Proposition 6, there exists *V*_1_ ⊆ *V*_0_ such that *E*_*s*_(*q*^*k*^) = *E*_*s*_(*V*_1_) and *E*_*t*_ (*q*^*k*^) = *E*_*t*_ (*V*_1_). Let *e*_*i*_ ∈ *E*_*s*_(*q*^*k*−1^) and *e*_*j*_ ∈ *E*_*t*_ (*q*^*k*−1^) be the two chosen edges in iteration *k*. Since 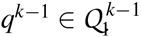, we know that either *e*_*i*_ ∈ *E*_*s*_(*V*_1_) and *e*_*j*_ ∈ *E*_*t*_ (*V*_1_), or *e*_*i*_ ∉*E*_*s*_(*V*_1_) and *e*_*j*_ ∉ *E*_*t*_ (*V*_1_). We further show the second case is not possible, since it immediately implies that *q*^*k*−1^ is not indivisible. Thus, we have that *e*_*i*_ ∈ *E*_*s*_(*V*_1_) and *e*_*j*_ ∈ *E*_*t*_ (*V*_1_). If *q*^*k*^ is obtained from the first situation (line 3) of Algorithm 3, then this already means that 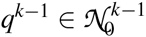. Otherwise (line 6), since we guarantee that the path p used in line 8 of Algorithm 3 does not contain any edge in *E*_*s*_(*q*^*k*−1^) ∪ *E*_*t*_ (*q*^*k*−1^), we can still conclude that 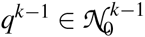, a contradiction, as desired.

### Proof of Proposition 9.

Since *q*^*k*^ is indivisible from Proposition 8, we know that during the *k*-th iteration, the two chosen edges *e*_*i*_ and *e*_*j*_ cannot have that *f* (*e*_*i*_) = *f* (*e*_*j*_), 1 ≤ *k* ≤ *n* − 1. According to Algorithm 3, we have that |*E*_*s*_(*q*^*k*^)| + |*E*_*t*_ (*q*^*k*^)| = |*E*_*s*_(*q*^*k*−1^)| + |*E*_*t*_ (*q*^*k*−1^)| − 1. After (*n* − 1) iterations, we must have that |*E*_*s*_(*q*^*k*−1^)| = |*E*_*t*_ (*q*^*k*−1^)| = 1, and both the two subsets become empty after the *n*-th iteration. Thus, after exactly (|*E*_*s*_(*q*)| + |*E*_*t*_ (*q*)| − 1) iterations, Algorithm 3 terminates.

### Proof of Proposition 10.

We first prove that, for 1 ≤ *k* ≤ *n* − 1, we have that Δ^*k*^ ≤ Δ^*k*-1^ Consider the change of number of edges and vertices during the *k*-th iteration. Let *e*_*i*_ and *e*_*j*_ be the two chosen edges. According to the proof of Proposition 9, we have that only *e*_*i*_ will be removed, and a new edge (namely, *e*_*k*_ in Algorithm 3) will be added. If it is the first situation (line 3 of Algorithm 3), then the middle vertex (namely, *v*_1_ = *u*_2_) cannot become isolated, since otherwise it implies that *q*^*k*−1^ is not indivisible. If it is the second situation (line 6 of Algorithm 3), some vertices in path *p* might be become isolated. But if this happens, then at least the same number of edges will also be removed in *p*. Thus, we always have that Δ^*k*^ ≤ Δ^*k*−1^.

We now show that Δ^*n*^ ≤ Δ^*n*−1^. This is because, in the *n*-th iteration, both *e*_*i*_ and *e*_*j*_ are removed, and only one edge will be added. Similar to the above analysis, if some vertices become isolated, then at least the same number of additional edges will be removed. Thus, we have that Δ^*n*^ ≤ Δ^*n*−1^ − 1.

According to our analysis in the main text, the merging process narrows down the solution space, i.e., we have |*P*^*k*∗^| ≥ |*P*^(*k*−1)∗^|, 1 ≤ *k* ≤ *n*. By combining these facts, we have that Δ^*n*^ − |*P*^*n*\*^| ≤ Δ − |*P*\*| − 1.

### Proof of Proposition 11.

The running time of Algorithm 4 is dominated by lines 2 and 3. According to Proposition 9, in Algorithm 3, Algorithm 6 will be called at most |*E* | times. Thus, according to Proposition 14 and the fact that 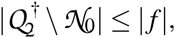, the total running time is *O*(|*V* |^2^ · |*E* |^2^ · | *f* |).

### Proof of Proposition 12.

Let (*P*, *w*) be any decomposition of (*G*, *f*). We can obtain a decomposition (*P*′, *w*′) of (*G*′, *f*) satisfying that |*P*′| = |*P*| and that *w*′ = *w* by only reversing the edges from *u* to *v* for each path in *P*. Since reverse operation is invertible, given a decomposition (*P*′, *w*′) of (*G*′, *f*), we can also obtain a decomposition (*P*, *w*) of (*G*, *f*) in the same way. This gives a one-to-one correspondence between the decompositions of (*G*, *f*) and that of (*G*′, *f*). And, in particular we have |*P*′^∗^| = |*P*^∗^|.

### Proof of Proposition 13.

The correctness of the algorithm is the consequence of the following facts. First, if 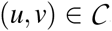, then among the vertices in *V* (*u*, *v*), *u* and *v* are the smallest and the largest vertices *w.r.t*. any topological ordering. This is simply because all vertices in *V* (*u*, *v*) \ {*u*, *v*} can be reached from u and can reach *v*. Second, *L* and *R* always store the current smallest and largest vertices in *S w.r.t.* the topological ordering. Thus, by definition, vertices added to *S* in line 9 of Algorithm 5 are guaranteed in the closure of *V*_1_. Third, *Q* always stores the vertices in *S* \ {*L*, *R*}. Thus, following the first fact, we know that vertices in *Q* cannot be the boundary vertices of the closure, and consequently, all their adjacent vertices (added in line 9) are guaranteed in the closure of *V*_1_. Fourth, when the algorithm terminates (line 14), we have that *S*= *V* (*L*, *R*). In fact, let *p* be any path from *L* to *R*. From line 13, we know that the first vertex in *p* is in *S*. Then following lines 8 to 11, all vertices in *p* will be added to *S*. Finally, all vertices in *S* \ {*L*, *R*} are examined such that all their adjacent vertices are in *S*. Thus, by definition, we know that (*L*, *R*) is closed. Besides, since all vertices added to S are guaranteed to in the closure of *V*_1_, we have that *S* is also the smallest subset, implying that (*L*, *R*) is the closure of *V*_1_. This reason also proves that the closure of *V*_1_ is unique. This algorithm is essentially the breadth-first search algorithm, in which edge each is examined at most once. Thus, its running time is *O*(|*E* |).

### Proof of Proposition 14.

If Algorithm 6 finds *w* ∈ (*S*_1_ ∩ *T*_2_) ∪ (*S*_2_ ∩ *T*_1_), then clearly we can perform a series of operations to make *e*_*i*_ and *e*_*j*_ adjacent. On the other side, suppose that there exists a series of reverse operations such that in the resulting graph we have that *e*_*i*_ = (*w*_1_, *w*) and *e*_*j*_ = (*w*, *w*_2_). Let *c*_1_ and *c*_2_ be the closures of {*w*_1_, *w*} and {*w*, *w*_2_}, respectively. Clearly, all the reverse operations that moves *e*_*i*_ (resp. *e*_*j*_) must operates on closures that are inside *c*_1_ (resp. *c*_2_). Thus, there must be a path from (*u*′_1_, *v*′_1_) (resp. (*u*′_2_, *v*′_2_)) to *c*_1_ (resp. *c*_2_) in 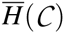, which implies that Algorithm 6 will have *w* ∈ (*S*_1_ ∩ *T*_2_) ∪ (*S*_2_ ∩ *T*_1_). This reasoning also implies that when Algorithm 6 traces back to recover the reverse operations (lines 8 and 9), the two paths are disjoint, since they are in two different subtrees (rooted at *c*_1_ and *c*_2_, respectively).

The running time of Algorithm 6 is dominated by building 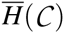, which is *O*(|*V* |^2^ · |*E* |).

